# Temporal and spatial dynamics in the apple flower microbiome in the presence of the phytopathogen *Erwinia amylovora*

**DOI:** 10.1101/2020.02.19.956078

**Authors:** Zhouqi Cui, Regan B. Huntley, Quan Zeng, Blaire Steven

**Author notes:** Corresponding author: Blaire Steven.; Quan Zeng.

## Abstract

Plant microbiomes have important roles in plant health and productivity. However, despite flowers being directly linked to reproductive outcomes, little is known about the microbiomes of flowers and their potential interaction with pathogen infection. Here, we investigated the temporal dynamics and spatial traits of the apple stigma microbiome when challenged with a phytopathogen *Erwinia amylovora*, the causal agent of fire blight disease. We profiled the microbiome from the stigmas of a single flower, greatly increasing the resolution at which we can characterize shifts in the composition of the microbiome. Individual flowers harbored unique microbiomes at the OTU level. However, taxonomic analysis of community succession showed a population gradually dominated by bacteria within the families *Enterobacteriaceae* and *Pseudomonadaceae*. Flowers inoculated *E. amylovora* established large populations of the phytopathogen, with pathogen specific gene counts of >3.0 × 10^7^ in 90% of the flowers. Yet, only 42% of inoculated flowers later developed fire blight symptoms. This reveals pathogen amount on the stigma is not sufficient to predict disease outcome. Our data demonstrate that apple flowers represent an excellent model in which to characterize how plant microbiomes establish, develop, and interact with biological processes such as disease progression in an experimentally tractable plant organ.

## Introduction

Flowers, the reproductive organs of angiosperms, play a critical role in the plant’s lifecycle. The most important function of flowers is to provide a mechanism for pollination, the union of sperm contained within pollen, to the ovules contained in the ovary. The fertilized ovules produce seeds that will later germinate to become the next generation of plants. Yet, unlike other vegetative organs such as the roots, stems, and leaves that are present through a large part of the plant’s lifecycle, flowers develop on mature plants and are typically present for the limited period during bloom. As such, research characterizing the microbiome of the flower is generally less developed than for other plant organs.

Flowers of apple (*Malus domestica*) have been subject to considerable research attention as they are the direct precursors of apple fruits, one of the most consumed fruits worldwide (1). The ephemeral nature of apple flowers, with mature flowers from petal open to petal fall only lasting for 5-10 days in spring, offers a unique environment in which to study community succession (1, 2). During bloom, petals open up in a relatively short period of time, typically within one day, which exposes the internal flower parts to the environment and microorganisms. Several of these internal flower parts exude various types of nutrient-rich secretions including nectar, stigmatic exudate, and pollen exudate, for the purpose of attracting pollinators, and inducing the germination of pollen grains (1, 3). These secretions are rich in sugars, amino acids, polysaccharides, and glycoproteins, which are excellent sources of nutrients for many microorganisms (1, 3, 4). The stigma is particularly nutrient rich and harbors a larger microbial biomass than other flower parts (5, 6). Previous research has documented a relatively low diversity of the stigma microbiome, although certain lineages predominantly within the families *Enterobacteriaceae* and *Pseudomonadaceae* tend to be dominant (7).

While the stigma provides an excellent niche for microbial colonization, it also offers an opportunity for pathogen infection. Many pathogens have evolved to take advantage of this environmental niche, among which one of the most important is the phytopathogenic bacterium *Erwinia amylovora*, the causal agent of fire blight. Fire blight is considered as one of the most devastating diseases of apple, with annual losses and costs of control estimated at over $100 million in the U.S. (8). During bloom, *E. amylovora* (*Ea*) cells are spread to apple flowers by insects, wind, or rain and multiply on the stigma surface (9). *Ea* cells can then migrate from the stigma to the hypanthium and enter into the host through the natural opening, the nectarthodes. Initial infection occurs at the ovary tissue and can spread to other parts of the plants through the plant vasculature system. Fire blight infection can result in significant yield reduction and / or tree death. In this regard, uncovering environmental or biologic factors that can inhibit the spread or development of fire blight are of considerable research interest.

One potential source of fire blight control is the natural microbiome of the stigma. Yet, there exist considerable knowledge gaps concerning how the stigma microbiome is established and structured. The studies that have considered the stigma microbiome have generally focused on cataloging microbial diversity through various culture-dependent and culture-independent methods (7, 10) and few studies have investigated the temporal development of the microbiome (2). Furthermore, previous research has predominantly studied the microbiome using pooled flower samples, thus it is uncertain the extent to which the microbiome differs among individual flowers of the same genetic background. Finally, how the colonization of a phytopathogen affects the development, composition, or structure of the stigma microbiome is essentially unknown. In this study, we examined the temporal development of the stigma microbiome in the presence and absence of *Ea* to investigate how this organism influences the development of the normal microflora of the apple flower stigma. Additionally, we characterized the variability of the microbiome amongst 100 individual stigmas inoculated with *Ea* to assess if certain microbiome members could regulate *Ea* colonization and growth on apple stigmas.

## Materials and methods

### Sampling site

To limit the effects of host and environmental conditions, we used flowers from nine trees of the same apple cultivar ‘Early Macoun’ (*Malus domestica* NY75414-1) planted at the same geographical location (Lockwood Farm, Hamden, Connecticut, 41.406 N 72.906 W). All trees were the same age and under the same maintenance program. Weather data (temperature and humidity) prior to and during bloom (from April 29^th^ to May 28^th^ 2018) is summarized in Table S1.

### Experiment design and stigma collection

#### Labeling flower clusters

On May 6^th^ 2018, 40 flower clusters that were in ‘King bloom’ stage (central flower opened but the four side flowers still closed, see Fig. 1A) were labeled with plastic tags. The day after the flower clusters were tagged, we identified clusters in which the side flowers were open and flower clusters with unopened flowers were not used. In this manner, only side flowers of roughly the same age were used in subsequent experiments.

**Figure 1.**
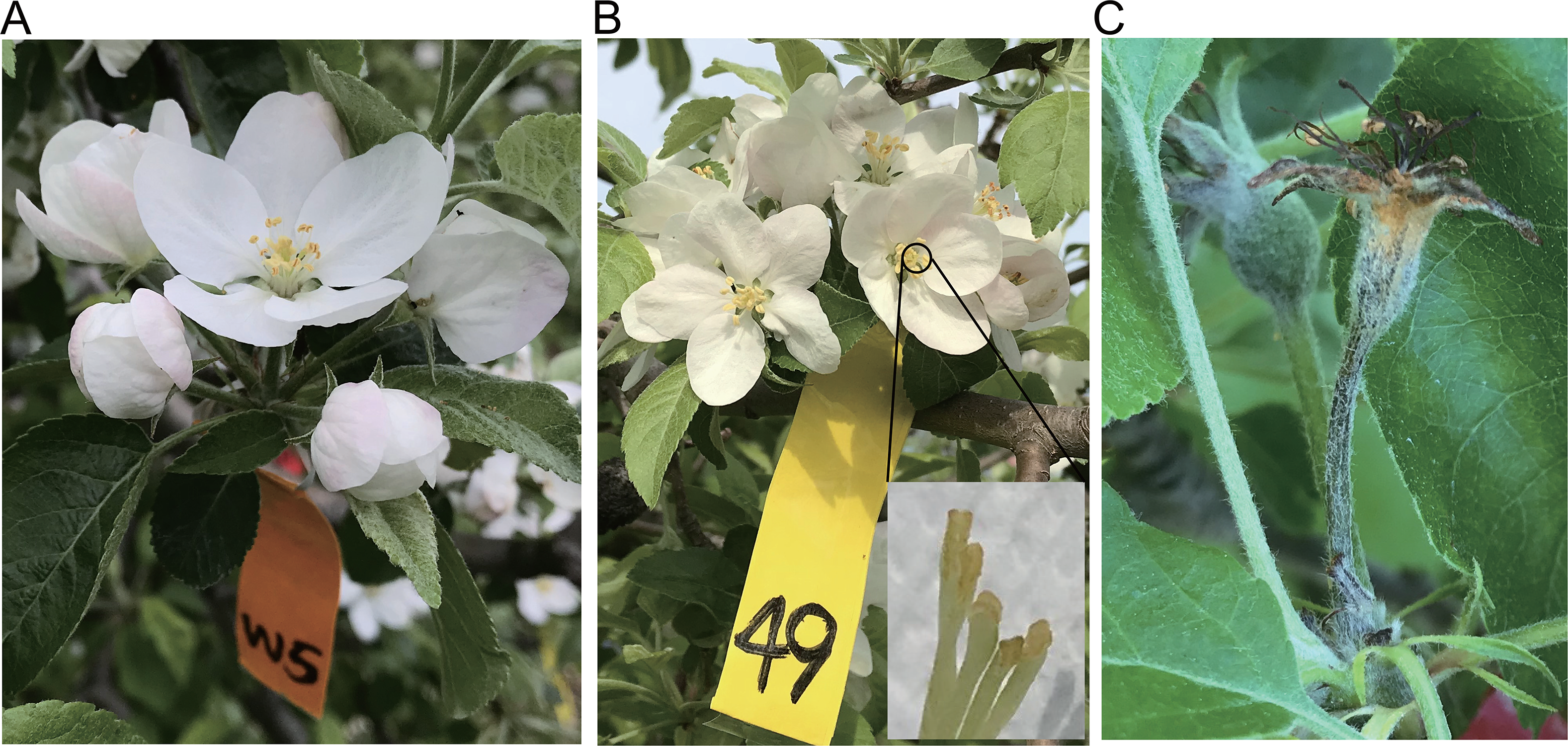
Illustration of apple flower clusters. (**A**) An apple flower cluster at “king bloom”. This is when flower clusters of the same age were tagged. (**B**) Once the surrounding flowers opened (one day after king bloom), stigmas of flowers were sampled and named as “day 1”. Each sample contains stigmas collected from an individual flower. A close up photo of individual stigmas is shown in the inset. (**C**) An example showing a flower with fire blight disease and a healthy flower coexisting in the same flower cluster. (Photo courtesy: Q. Zeng)

#### Sampling for temporal alterations of the stigma microbiome

On May 7^th^ 2018, ten of the 40 tagged flower clusters were selected, and the stigmas of an individual flower were harvested with sterile scissors (see Fig. 1B) and placed in a sterile 1.5 ml microcentrifuge tube. Collected stigma samples were kept in liquid nitrogen and transported to the laboratory for immediate processing. These samples were labeled as day 1 samples. The next day, another 10 flowers were selected for DNA extraction as described above (as day 2 samples). Immediately after sample collection on day 2, *Ea* was inoculated onto 20 tagged flower clusters and labeled as *Ea* treated. The inoculum consisted of an overnight culture of *E. amylovora* 110 grown in lysogeny broth (LB) diluted to a final concentration of 1 × 10^6^ CFU ml^−1^ in sterile water. The diluted culture was spray-inoculated to the open flowers using a handheld sprayer to ensure every flower was evenly exposed. Another twenty flower clusters were sprayed with sterile water as water controls. On each subsequent day (day 3 to day 5), stigmas from 20 *Ea*-treated and 20 water-treated flowers were collected and processed according to the same method described above.

#### Sampling for spatial patterns in the stigma microbiome

To investigate a larger spatial sampling of *Ea* inoculated flowers, we performed a parallel experiment, and tagged an additional 150 flower clusters to ensure flowers used in the experiment were the same age as the rest of the experimental set. As described for the temporal sampling, the flower clusters were individually spray-inoculated with *Ea* (1 × 10^6^ CFU ml^−1^) on day 2, and stigma samples of individual flowers were harvested on day 4. A total of 100 flowers of the same developmental stage were harvested for DNA extraction, while the remaining flowers of each flower cluster were left on the tree to monitor disease development. Blossom blight symptoms, black withering and dying of the remaining flowers (Fig. 1C), were evaluated two weeks after inoculation on May 24^th^, 2018. An illustrated scheme of both temporal and spatial sampling is shown in Fig. S1.

#### DNA extraction and sequencing of bacterial 16S rRNA genes

For extraction of bacterial DNA, 200 μl of 0.5x phosphate-buffered saline (PBS) was added to each microcentrifuge tube containing stigma samples. Epiphytic microbes were removed from the stigma by a 5-minute water bath sonication followed by a 30-second vortex. DNA was extracted from the 200 μl of bacterial suspension by using the DNeasy PowerSoil Kit (Qiagen, Hilden, Germany) according to manufacturer’s instructions. The amount of template DNA added in the PCR reaction (25 μl) ranged from 10.0 ng to 20.0 ng as determined by Nanodrop2000 (Thermo Fisher Scientific, Waltham, MA). DNA was amplified by using the 515f/806r primer set, which targets the V4 region of the bacterial 16S rRNA gene, with both primers containing a 6-bp barcode unique to each sample (11). PNA clamps were added to the PCR mixture at a concentration of 0.75 μM to block the PCR amplification of apple plastid and mitochondrial sequences (7). PCR conditions were performed as described in Steven et al. (2018) (7). Successful PCR amplifications at the correct amplicon size were confirmed by gel electrophoresis. The PCR products were purified and normalized by using SequalPrep normalization plate kit (Invitrogen, CA, USA). Pyrosequencing was conducted on an Illumina MiSeq v2.2.0 platform through services provided by the UConn MARS facility.

#### Quantitative PCR for enumeration of E. amylovora

The abundance of *Ea* in each collected stigma sample was quantified by determining the cycle threshold (CT) value of the *Ea* specific gene *amsC* (12). Quantitative PCR (qPCR) was performed using a SsoAdvanced universal SYBR Green supermix (Bio-Rad, CA, USA), as described previously (13). The CT values for a 1/10 dilution series of known *amsC* gene copies of *E. amylovora* chromosomal DNA was determined to make a standard curve for calculation of copy numbers in stigma samples.

### Bioinformatics and statistical analysis

Illumina sequencing reads were assembled into contigs and quality screened by using mothur v1.39.5 as previously described (14). Sequences that were at least 253 bp in length, contained no ambiguous bases, and no homopolymers of more than 8 bp were used in the analysis. Chimeric sequences were identified by using the VSEARCH as implemented in mothur (15), and all potentially chimeric sequences were removed. To maintain a similar sampling effort between samples, samples with less than 10,000 sequences per sample were also removed. The resulting sequence counts per sample are presented in Table S2. Negative control (PCR using sterile H_2_O as a template) was also included in both sequence datasets. The sequences data are deposited in the Sequence Read Archive under accession number PRJNA597302.

Sampling effort was normalized to the depth of the smallest sample and operational taxonomic units (OTUs) were defined at 100% sequence identity, employing the OptiClust algorithm in mothur (16). Taxonomic classification of sequences was performed with the Ribosomal Database Project (RDP) classifier against the SILVA v132 reference alignment in mothur (17, 18). Non-metric multidimensional scaling (NMDS) was used to visualize the pairwise distances among samples with Bray-Curtis distances in the Vegan package in R (19). Descriptive diversity statistics were calculated in mothur. The correlation between alpha diversity determined with the non-parametric Shannon’s Diversity Index and *E. amylovora* abundance in each sample was generated with the ggplot2.0 package for R (20). Statistically significant differences in diversity statistics were identified with a one-way ANOVA and Tukey-Kramer post hoc test in the agricolae package in R.

## Results

### Temporal patterns in stigma microbial community assembly

We characterized the microbial community on stigmas collected from individual flowers, over a period of 5 days after petal opening, to investigate the temporal dynamics in community assembly and microbial succession on the stigma. Meanwhile, we included *Ea* inoculated stigmas to compare community succession in the presence of a phytopathogen. A total of 2 930 231 high-quality filtered sequences were obtained from 96 samples with the number of sequences ranging from 10 210 to 97 668 (Table S2). These sequences clustered into 46 809 OTUs (mean 222 per sample) at 100% sequence similarity.

At the phylum level, 24 phyla were detected. In both the water control and *Ea* inoculated datasets, the dominant phylum was *Proteobacteria* (94.3% of total sequences), followed by *Cyanobacteria* (3.6%), *Actinobacteria* (0.8%), *Firmicutes* (0.2%) and *Bacteroidetes* (0.2%). A temporal pattern was observed, in that phyla outside the *Proteobacteria* were most abundant in the early time points (days 1 and 2) accounting for 15% of sequences and decreasing to <1% at later time points (Fig. S2).

Given the dominance of *Proteobacteria*, these sequences were classified to deeper taxonomic ranks. Sixty-seven families were identified, with the majority belonging to the *Enterobacteriaceae* (average 70.0%, blue bars) and *Pseudomonadaceae* (26.2%, red bars in Fig. 2), with small contributions from *Moraxellaceae* (0.6%)*, Beijerinckiaceae* (0.2%), unclassified *Gammaproteobacteria* (0.3%)*, Burkholderiaceae* (0.3%) and *Xanthomonadaceae* (0.2%) (Fig. 2). Of note, both *Pseudomonadaceae* and *Enterobacteriaceae* gradually accounted for a larger proportion of the microbiome as time progressed in both water control and *Ea* inoculated datasets (Fig. 2). Yet, the average proportion of *Enterobacteriaceae* (the family to which *Ea* belongs) was higher in the *Ea* treated flowers compared to water control (89.7% versus 45.6% at day 5) (Fig. 2).

**Figure 2.**
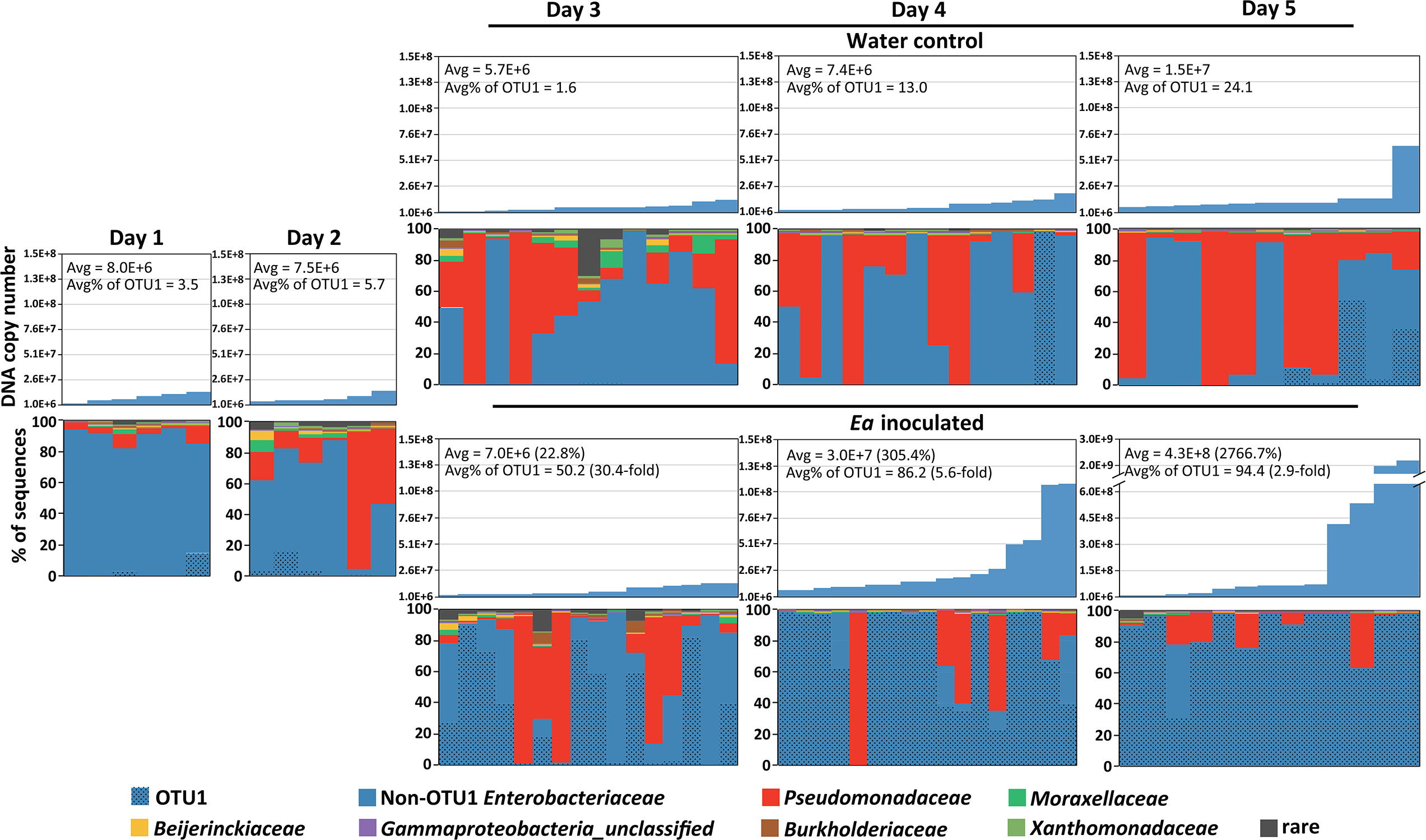
Temporal dynamics in the predominant bacterial families present on stigmas of individual flowers. Each column represents a single flower. The seven most abundant families are displayed, and the category “rare” represents the sum of the remaining taxa. The relative abundance of OTU1, identified as sharing 100% sequence identity with *Erwinia amylovora*, is indicated by hatched lines. Copy numbers of *E. amylovora amsC* gene in each sample were determined by qPCR, and are displayed in the bar graphs above the stacked columns. The average DNA copies are indicated as well as the average relative abundance of OTU1. The change in *Ea* inoculated compared to water control was labeled in the brackets. Water control: flower clusters sprayed with sterile H_2_O. *Ea* inoculated: flower clusters sprayed with a bacterial suspension of *E. amylovora* strain 110. Day 1-day 5 represent the number of days after petals opened during bloom.

### Abundance of Ea on individual flowers

We employed two methods to assess the abundance of *Ea* in the datasets, relative abundance of *Ea* sequences in the dataset and *Ea* copy numbers quantified by qPCR of an *Ea* specific gene. First, we identified an OTU in the dataset that had 100% sequence identity with the inoculated *Ea* strain (OTU1; Table S3). OTU1 was detected every day but not in all samples. On days 1 and 2, prior to the stigma treatments, OTU1 made up an average of 4% and 6% of the microbiome sequences, respectively (filled bars in Fig. 2). In the control water sprayed stigmas the proportion of OTU1 gradually increased from an average of 2% of sequences on day 3 to 13% on day 4, finally making up an average of 24% of sequence on day 5. In contrast, the populations of OTU1 were larger in the *Ea* treated stigmas. By day 3 OTU1 accounted for an average of 50% of the sequence libraries, increasing to 86% on day 4 and ending at 94% of sequences on day 5, a 2.9-fold increase in comparison to the controls (Fig. 2).

In addition, qPCR was performed to quantify the genome copies of *Ea* in each stigma sample. As was observed for OTU1, *Ea* was identified across the dataset. In the pretreated stigmas (days 1 and 2) the average copy number of *Ea* DNA were ~7.7 × 10^6^ (Fig. 2). In the control datasets, the DNA copy number were similar on days 3 and 4 at 5.7 × 10^6^ and 7.4 × 10^6^, respectively, and increased to 1.5 × 10^7^ on day 5 (Fig. 2). In the *Ea* inoculated flowers the copy number of *Ea* on day 3 (one day after inoculation) was similar to the control flowers, suggesting *Ea* had not yet established strong growth on the stigma (Fig. 2). However, by day 4 the average abundance of *Ea* on the treated stigmas reached 3.0 × 10^7^, a 300% increase compared to the controls, and increased further on day 5 reaching an average of 4.3 × 10^8^, a 28-fold increase in comparison to the controls (Fig. 2). Taken together, these data suggest that *Ea* may be naturally present in the orchard, as it was commonly detected in the pretreated and control stigmas. Yet, the *Ea* inoculation clearly benefited *Ea* colonization, which was readily apparent by day 5, three days after the inoculation.

### Effects of Ea inoculation on community composition and diversity

To test if *Ea* treatment had a significant effect on microbiome composition, we visualized the Bray-Curtis distances among samples of each dataset using NMDS. The samples clearly clustered due to *Ea* inoculation, which was confirmed by permutational multivariate ANOVA (P = 0.001) (Fig. 3A). Additionally, samples were also clustered based on days post-bloom (P = 0.001) (Fig. 3A). Diversity of the stigma communities was assessed by calculating the Shannon’s Diversity index. For both control and *Ea* inoculated datasets there was a trend towards increased diversity in the early time points, which then decreased by days 4 and 5 (Fig. 3B). When the control and *Ea* inoculated datasets were combined to test the overall effect of pathogen presence on microbial diversity, there was no significant difference in diversity due to *Ea* treatment (p=0.109; Fig. 3B).

**Figure 3.**
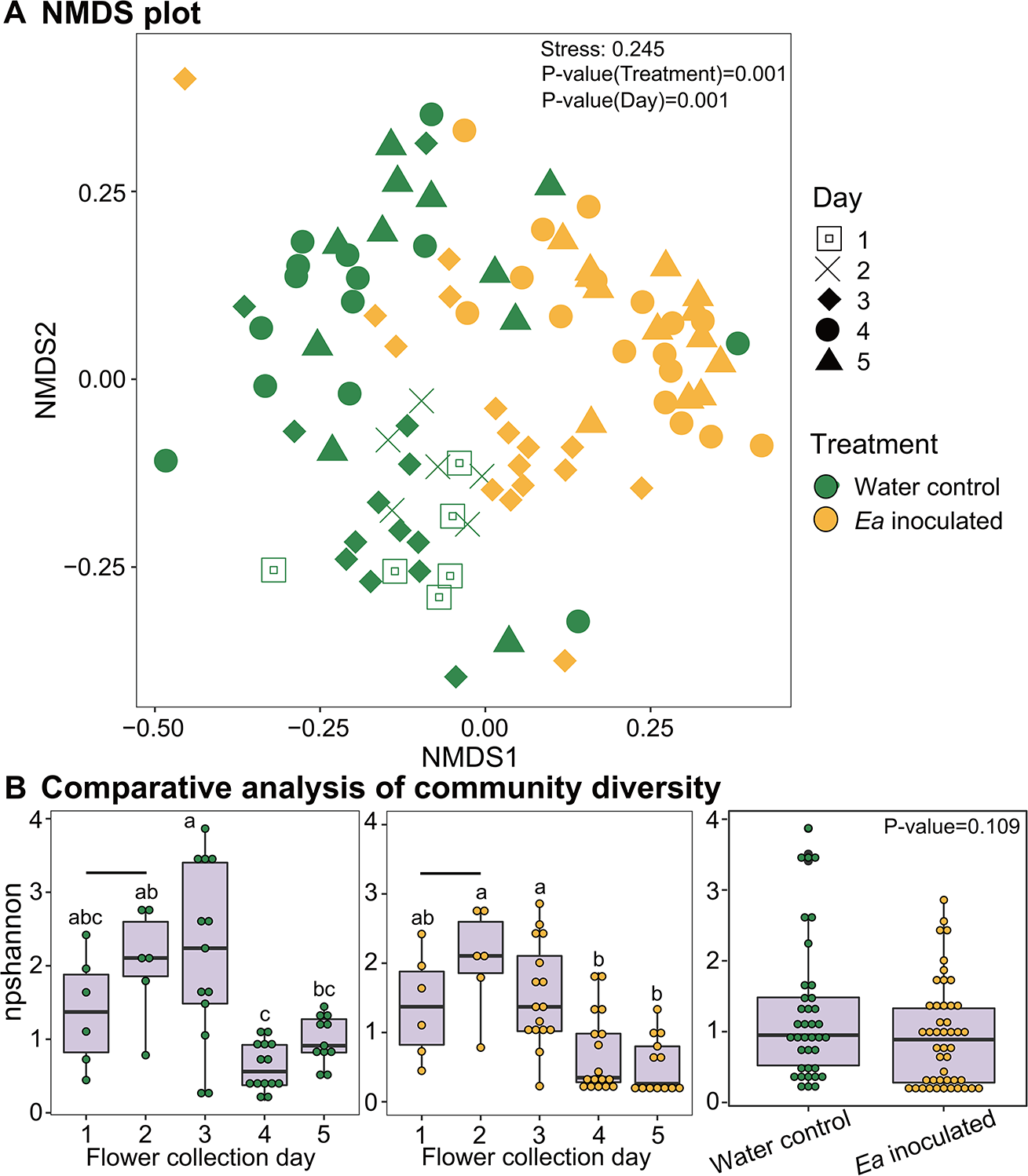
(**A**) Non-metric Multidimensional Scaling (NMDS) plot displaying relationships of stigma microbial community composition in samples from water control (green) and *Ea* inoculated samples (gold). Symbols indicate stigma sample collection day. The distances were determined using the Bray-Curtis metric and the stress value of the ordination is indicated. Statistically significant differences in clustering were evaluated via the Adonis permutation test and P-values are indicated. (**B**) Comparative analysis of community diversity (Shannon index) among stigma samples. Changes in diversity over time for the water control samples (left panel) and the *Ea* inoculated samples (middle panel), respectively. The bar above day 1 and day 2 indicates the pre-treatment samples, which are the same between the panels. Overall diversity of water control samples versus *Ea* inoculated samples (far right panel). Statistically significant differences were identified by ANOVA comparisons of means, employing a post-hoc Tukey-Cramer test for multiple comparisons. Boxes labeled with different letters showed statistically significant differences (P-value <0.05).

Collectively, these findings indicate that taxonomically diverse microbial populations initially colonize the stigma of the apple flower. Gradually, a community dominated by representatives of the *Pseudomonadaceae* and *Enterobacteriaceae* families outcompetes these populations and become the predominant community members (Fig. 2), which results in an overall decrease in diversity of the stigma microbial community (Fig. 3B &C). In the face of *Ea* challenge there is a significant shift in the composition of the microbial community (Fig. 3A). Yet, there is no significant effect on the diversity of the community as a whole in comparison to the control flowers (Fig. 3B).

### The influence of Ea inoculation on 100 spatially separated flower clusters

The data for the temporal dynamics were based on a limited number of samples. To further explore the interaction of microbes when colonized by a phytopathogen, we expanded the analysis to 100 spatially separated flower clusters (approximately 400 individual flowers) inoculated with *Ea*. Flowers were collected from the clusters for microbiome characterization, while the remainder of the flowers were left on the tree to monitor the rate of disease development. Three weeks after *Ea* inoculation, only 42.4% of the flowers developed fire blight symptoms. Given that the genetic background of the host, flower age, and pathogen exposure were all identical between the inoculated flowers, and the trials were all performed in the same orchard and thus under the same environmental conditions, these observations suggest that none of these factors are sufficient to explain or predict disease occurrence at the single flower level.

### Genome copies of Erwinia amylovora

We measured the *amsC* copy number from 100 individual flowers by qPCR. The copy number varied from 1.3 × 10^4^ to 3.7 × 10^10^. The average was 4.4 × 10^9^ (dashed line, Fig. 4A) and the majority (90%) of stigmas harbored > 3.0 × 10^7^ gene copies of *Ea*, which is similar to the average of day 4 inoculated flowers in the temporal dynamics study. These results indicate most of the stigmas harbored large populations of *Ea*, despite only a proportion of flowers later developing fire blight symptoms.

**Figure 4.**
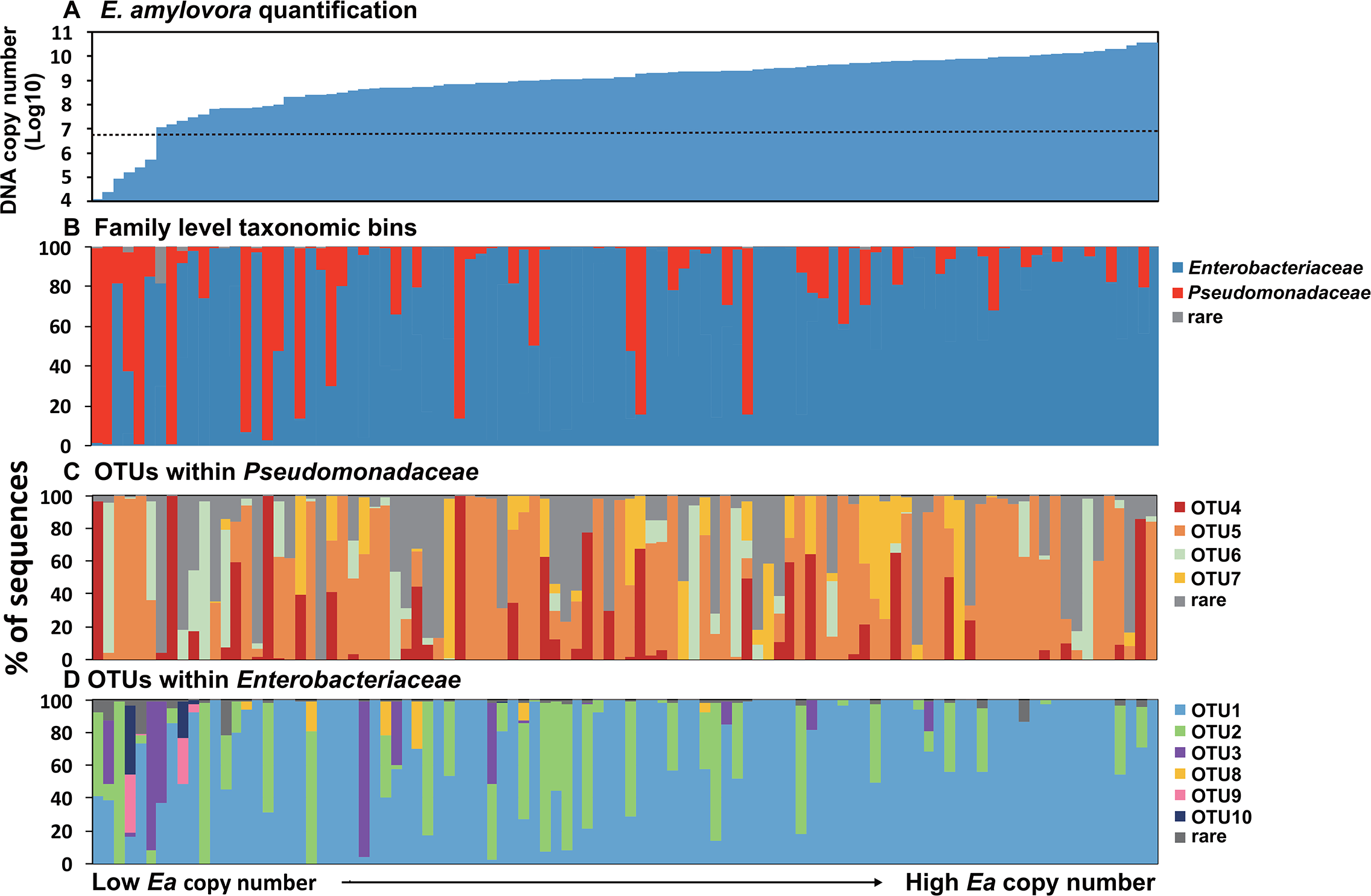
(**A**) DNA copy numbers of the *Ea* specific gene *amsC* ordered by abundance in 100 flowers. The dashed line represents average copy number across the samples. (**B**) Relative abundance (%) of the two major bacterial families within *Proteobacteria* in the stigma microbiome of 100 flowers. Each column represents an individual flower. The columns are ordered by *amsC* copy number to match Fig. 4A. (**C**) OTUs within the family *Pseudomonadaceae* and (**D**) *Enterobacteriaceae.* The category “rare” represents the sum of the remaining taxa.

### Microbiome composition

A total of 4 176 840 high-quality 16S rRNA gene sequences were recovered from the 100 flowers, with the number of sequences ranging from 19 297 to 80 130 per sample. After normalizing sampling to the smallest dataset, clustering produced 27 843 OTUs (mean 282 per sample) at 100% sequence similarity. The detailed information for each dataset is presented in Table S2.

At the phylum level, 22 phyla were identified among the sequences. The most abundant, *Proteobacteria*, ranged from 96.8% to 100% of recovered sequences, followed by *Actinobacteria* (0-0.5%), *Cyanobacteria* (0-1.6%) and *Firmicutes* (0-1.5%) (Fig. S3). Within the *Proteobacteria*, 59 families were identified and *Pseudomonadaceae* (red bar in Fig. 4B) and *Enterobacteriaceae* (blue bar) were predominant (> 81.5% in each sample). Notably, the proportion of *Pseudomonadaceae* and *Enterobacteriaceae* significantly varied among the 100 samples, from 0.02% to 99.20% and from 0.45% to 99.97%, respectively (Fig. 4B).

### OTUs within the Pseudomonadaceae and Enterobacteriaceae

Sequences within *Pseudomonadaceae* and *Enterobacteriaceae* were classified to deeper taxonomic ranks to investigate if particular OTUs were associated with *Ea* abundance. Of the 10 most abundant OTUs in the dataset, four belonged to the *Pseudomonadaceae* and six to the *Enterobacteriaceae*, representing five different genera (Table S3). By in large each flower harbored a unique microbiome composition, with widely varying abundance of the predominant OTUs among the samples (Fig. 4C &D). Furthermore, there was no observable pattern in specific OTUs being co-abundant in the samples with a high relative abundance of *Pseudomonadaceae* (Fig. 4B &C). For example, when we tested the correlation between the relative abundance of the most abundant *Pseudomonadaceae*-related OTU (OTU5; Table S3) and the relative abundance of *Pseudomonadaceae* in the dataset, the result showed no relationship (R^2^ = 0.26, Fig. S4). In other words, it was not a specific OTU that accounted for the high prevalence of the family *Pseudomonadaceae*. In contrast, OTU1 (100% sequence identity to *E. amylovora*; Table S3) tended to be highly abundant in samples with elevated counts of *Ea* (Fig. 4A &D). However, in those samples with low *Ea* counts a particular *Enterobacteriaceae* OTU was not predominant, suggesting that a specific OTU was not outcompeting *Ea* in those samples in which *Ea* was not well established.

### Correlates of Ea abundance to metrics of the stigma microbiome

To test if there were any aspects in the community data that were predictive of *Ea* abundance we performed four correlational analyses. First, the most abundant OTU in the dataset (OTU1) shared 100% sequence identity with the inoculated *Ea* strain (Table S3). Therefore, we tested the correlation between the relative abundance of OTU1 and the *amsC* gene copy number of *Ea*, and thereby testing if the relative abundance of OTU1 was correlated to *Ea* absolute abundance (Fig. 5A). The result showed there was a positive relationship between the two metrics, with an R^2^ = 0.29, suggesting a relationship but low explanatory power. Second, as shown in Fig. 4B, many of the stigmas maintained a large proportion of *Pseudomonadaceae* populations. We investigated if there was a predictive relationship between the relative abundance of the *Pseudomonadaceae* and the copy number of *Ea.* The relationship displayed in a negative pattern but again had a low predictive value (R^2^ = 0.26, Fig. 5B). Thus, an increasing proportion of *Pseudomonadaceae* was not associated with a reduction of *Ea* colonization or abundance. Finally, we tested if *Ea* abundance was correlated to two different metrics of diversity of the stigma microbiome, Shannon’s diversity index and the number of recovered OTUs. In both cases there was no relationship between *Ea* abundance and diversity (Fig. 5C &D). Thus, there was no apparent effect of *Ea* abundance on the overall diversity of the stigma microbial communities.

**Figure 5.**
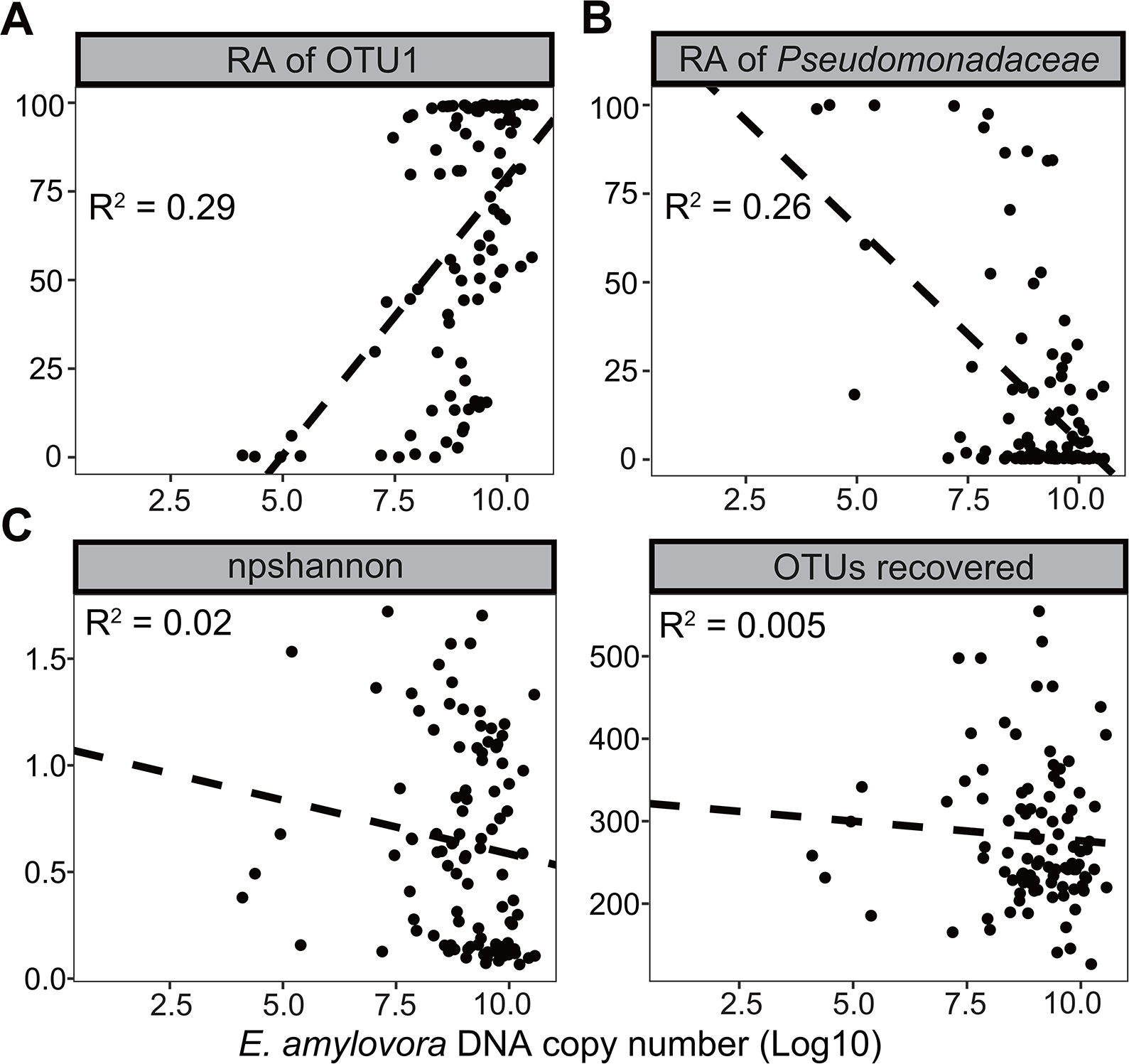
Correlations of *Ea* abundance as measured by qPCR against metrics of microbiome composition. (**A**) Relative abundance (%) of OTU1 identified as *Ea* (*R^2^* = 0.29, *P* = 0.28), (**B**) Relative abundance (%) of sequences within the *Pseudomonadaceae* family (*R^2^* = 0.26, *P* = 0.00), (**C**) community diversity (Shannon index) (*R^2^* = 0.02, *P* = 0.19), and (**D**) Number of recovered OTUs (*R^2^* = 0.005, *P* = 0.49). The dashed line is best fit from a linear model test. RA: relative abundance.

## Discussion

The apple flower microbiome has been previously recognized as an important factor for plant health and as a potential source of biocontrol agents against plant pathogens (1, 10, 21). Additionally, since the stigma is the major site of pollination and supports the growth of a large microbial population, microbial growth on the stigma may also influence pollination (22, 23). Thus, the stigma of a flower is a particularly important plant tissue for studying the microflora that associate with plants. Yet, information concerning the establishment, composition, and development of the microbiome on flower stigmas, as well as the disturbance by the colonization of a phytopathogen, are largely lacking. Previous studies have generally described the flower microbiome from whole flowers or nectar (1, 2, 24). In this study, we present data based on collecting the stigmas from a single flower, increasing both the temporal and spatial resolution at which the microbiome can be characterized.

Temporal dynamics are important for understanding the evolution of microbial communities (25–27). Shade et al. (2013) characterized the development of the microbiome on pools of apple flowers under a management program of treating the flowers with the antibiotic streptomycin to control fire blight. They found bacteria in the phyla TM7 and *Deinococcus* were predominant and showed signals of ecological successions with flower age (2). In our study, bacteria within the families *Pseudomonadaceae* and *Enterobacteriaceae* were numerically dominant (Fig. 2), which is more congruous with other studies of both the culture-dependent (10) and culture-independent characterizations (7) of apple flower microbial populations. This discrepancy is likely due to methodological differences between studies, or PCR biases induced by different PCR primer and blocking pairs. In either case, both studies identified strong signals of temporal patterns in how the microbiome is structured with flower age. The data presented here points to a core microbiome that was gradually established on the stigma predominantly composed of *Pseudomonadaceae* and/or *Enterobacteriaceae* within the phylum *Proteobacteria* (Fig. 2). The succession of these families was associated with a reduction of other bacterial taxa, such as the *Moraxellaceae, Xanthomonadaceae,* and *Burkholderiaceae,* which were only present in the early stages of bloom (Fig. 2). Concurrently, the later stages of bloom were associated with a lower diversity, supporting the observation that a small number of taxa had monopolized the stigma environment as the flower aged (Fig. 2). These observations are consistent with the stigmas being open to colonization by numerous bacteria in the initial stages of bloom. As the petals open, multiple bacteria carried by wind, dew or insects are introduced to the stigma creating a diverse microbial population (9). However, with time those bacteria best adapted to the stigma environment prevail and flourish. This is analogous to other observations, that complex microbial communities inoculated into a simple medium converge on a state similarly composed of bacteria in the families *Pseudomonadaceae* and *Enterobacteriaceae,* a phenomenon referred to as “emergent simplicity” (28). Thus, there may be conserved rules that govern the assembly of microbial communities, with respect to niche adaptation (5, 6), and microbial competition (29). Yet, predicting specific microbiome states of individuals or whether the factors that govern community assembly are deterministic or stochastic still remain significant knowledge gaps.

Inoculation of the flowers with *Ea* induced a significant shift in the structure of the microbiome (Fig. 3A). The data indicated that the abundance of *Ea* did not alter microbiome diversity (Fig. 5C &D), but *Ea* abundance may be negatively correlated with the presence of other microbes, particularly within the family *Pseudomonadaceae* (Fig. 5B). Most notably 90% of inoculated flowers inhabited large counts of *Ea* (> 3.0 × 10^7^ gene copies) and a high relative abundance of sequences identical to the inoculated pathogen (Fig. 4A & D), yet less than half of the flowers (42%) later developed fire blight symptoms. As the stigma sampling for microbiome characterization is necessity destructive, we cannot definitively link the status of the microbiome to disease development. However, these data strongly point to the absolute abundance of *Ea* to be a poor predictive measurement of disease occurrence. Thus, there must be another bottleneck in fire blight disease development beyond *Ea* growth on the stigma. These could include microclimate (30), antimicrobial compounds or yeasts in the nectar (10, 24), and host system sensing signals of high bacterial density (31). Yet, the observation of a high carrier rate of a pathogen with low disease incidence is synonymous with reports for many human pathogens. For instance, it is well established that 20-40% of the population are asymptomatic persistent carriers of *Staphylococcus aureus,* with a further 70-90% of people considered transient carriers (32). Yet only a minority of people will develop diseases such as sepsis, pneumonia, or osteomyelitis caused by *S. auerus* infection (33, 34). Similar phenomenon are observed for *Cutibacterium acnes* as a contributor to skin acne, which is also a major population in the healthy skin microbiome (35). In this respect, the dynamics of *Ea* growth and fire blight development appear to follow similar dynamics of other diseases, with a high carrier rate, but lower disease incidence.

## Conclusion

In this study we show that the apple flower stigma microbiome shares many characteristics with other host microbiome systems. In the initial stages of stigma colonization, the microbiome is temporally dynamic, which eventually settles into an equilibrium community (Fig. 2). Similar dynamics have been found in mammalian infants, fish, and soil (36-38). At the OTU level, individual flowers harbor largely unique microbiomes (Fig. 4C &D), similar to vertebrates and insects (39-41). Despite the diversity of the stigma microbiome at the OTU level (~200 OTUs per sample), the OTUs fell into just two predominant families (*Pseudomonadaceae* and *Enterobacteriaceae*) that differed in abundance between individual flowers (Fig. 2 &4). This mirrors the observation of the dominance of the *Firmicutes* and *Bacteroidetes* in the human intestinal tract, the so-called *Firmicutes/Bacteroidetes* ratio, and its potential influence on characteristics such as obesity (42, 43). Finally, we observe that virtually all flowers exposed to the phytopathogen *E. amylovora* developed large pathogen loads (Fig. 4), yet only a fraction (~42%) of the flowers developed disease, reflecting a common observation that pathogen burden is not always predictive of disease development (31, 44). Thus, we propose that the stigma microbiome is not only an important system to potentially identify biocontrol agents for impeding the development of fire blight, but represents a model system that can be employed to investigate the rules that govern microbial community assembly, development, and influence on disease progression and severity.

## Acknowledgments

We thank Sali Diallo and Zach Seltzer for their technical support. This study was supported by the USDA-NIFA-Organic Transitions grant 2017-51106-27001, Northeastern IPM Center partnership grant and USDA-Specialty Crop Block Grant (SCBG) through the Department of Agriculture, State of Connecticut.

Supplementary information is available at ISME’s website.

---

**Figure S1**. Schematic diagram describing temporal dynamics and spatial distribution sampling.

**Figure S2.** Temporal dynamics in the predominant bacterial phyla present on stigmas of individual flowers. Each column represents a single flower and are ordered by *Ea* abundance as determined by qPCR to match Fig. 2. The five most abundant phyla are displayed, and the category “rare” represents the sum of the remaining taxa. Water control: flower clusters sprayed with sterile H_2_O. *Ea* inoculated: flower clusters sprayed with a bacterial suspension of *E. amylovora* strain 110.

**Figure S3.** Relative abundance (%) of the four major bacterial phyla in the stigma microbiome of 100 flowers. Each column is an individual flower and are ordered by *amsC* copy number to match Fig. 4A. The five most abundant phyla are displayed, and the category “rare” represents the sum of the remaining taxa.

**Figure S4.** Correlational analysis in the relative abundance of OTU5 (most abundant OTU within the *Pseudomonadaceae*) against the relative abundance of *Pseudomonadaceae* (*R^2^* = 0.26, *P* = 0.00). The dashed line is best fit from a linear model test.

