## Supplementary figures and images for "Temporal and spatial dynamics in the apple flower microbiome in the presence of the phytopathogen *Erwinia amylovora*"

### Fig S1

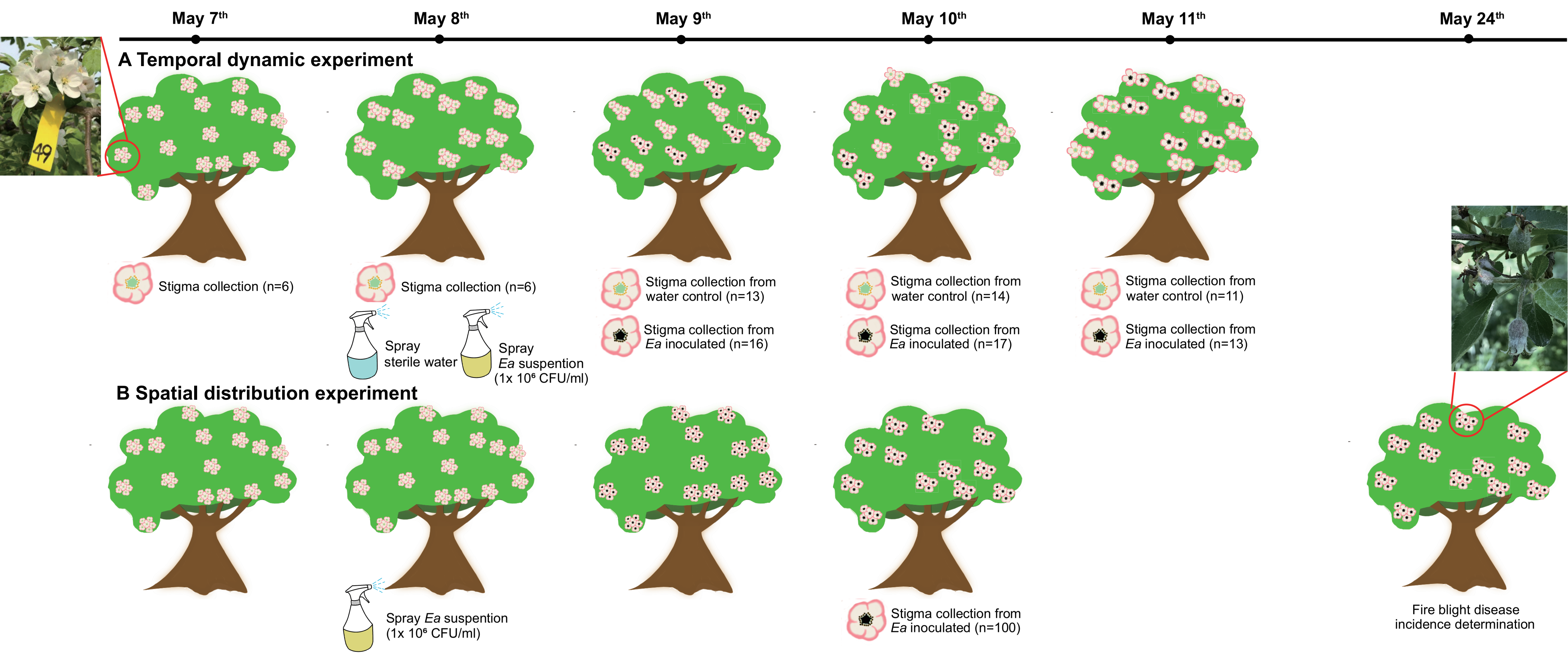

### Fig S2

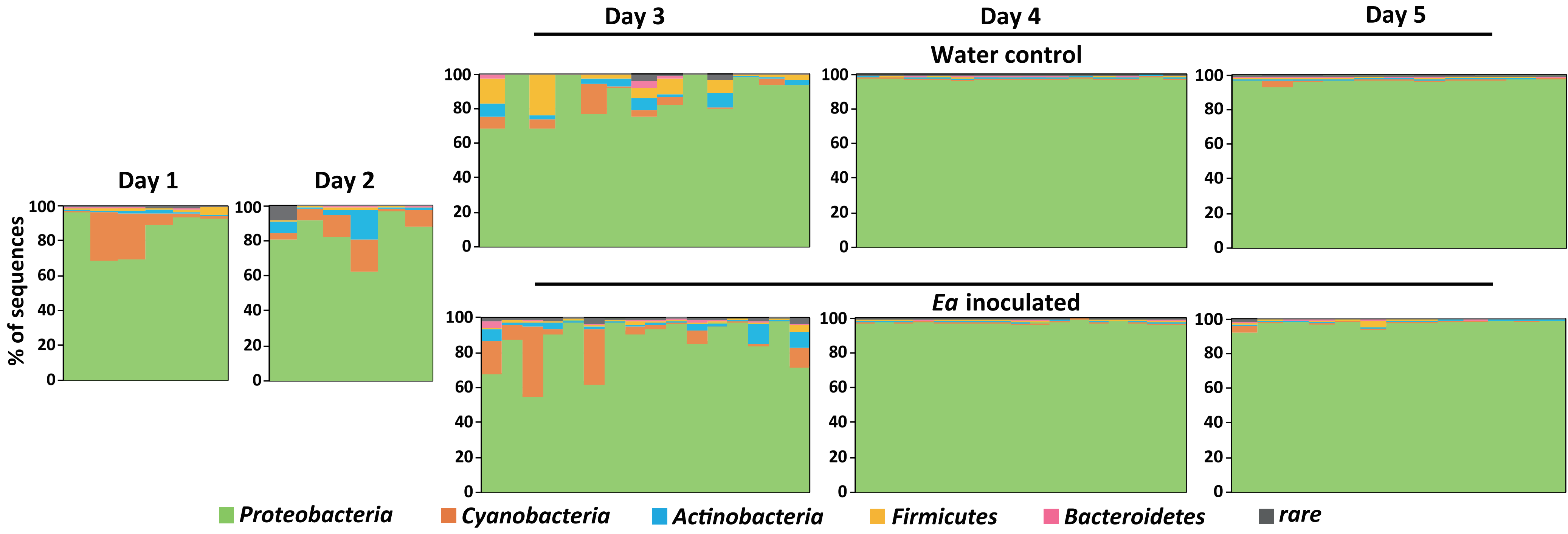

### Fig S3

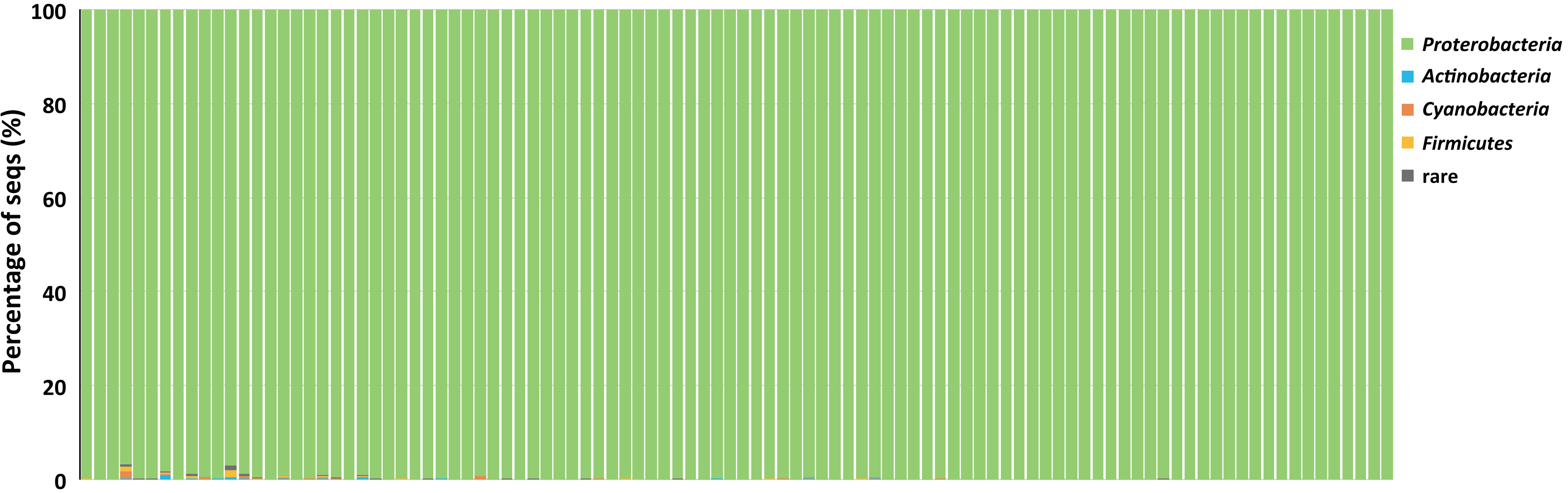

### Fig S4

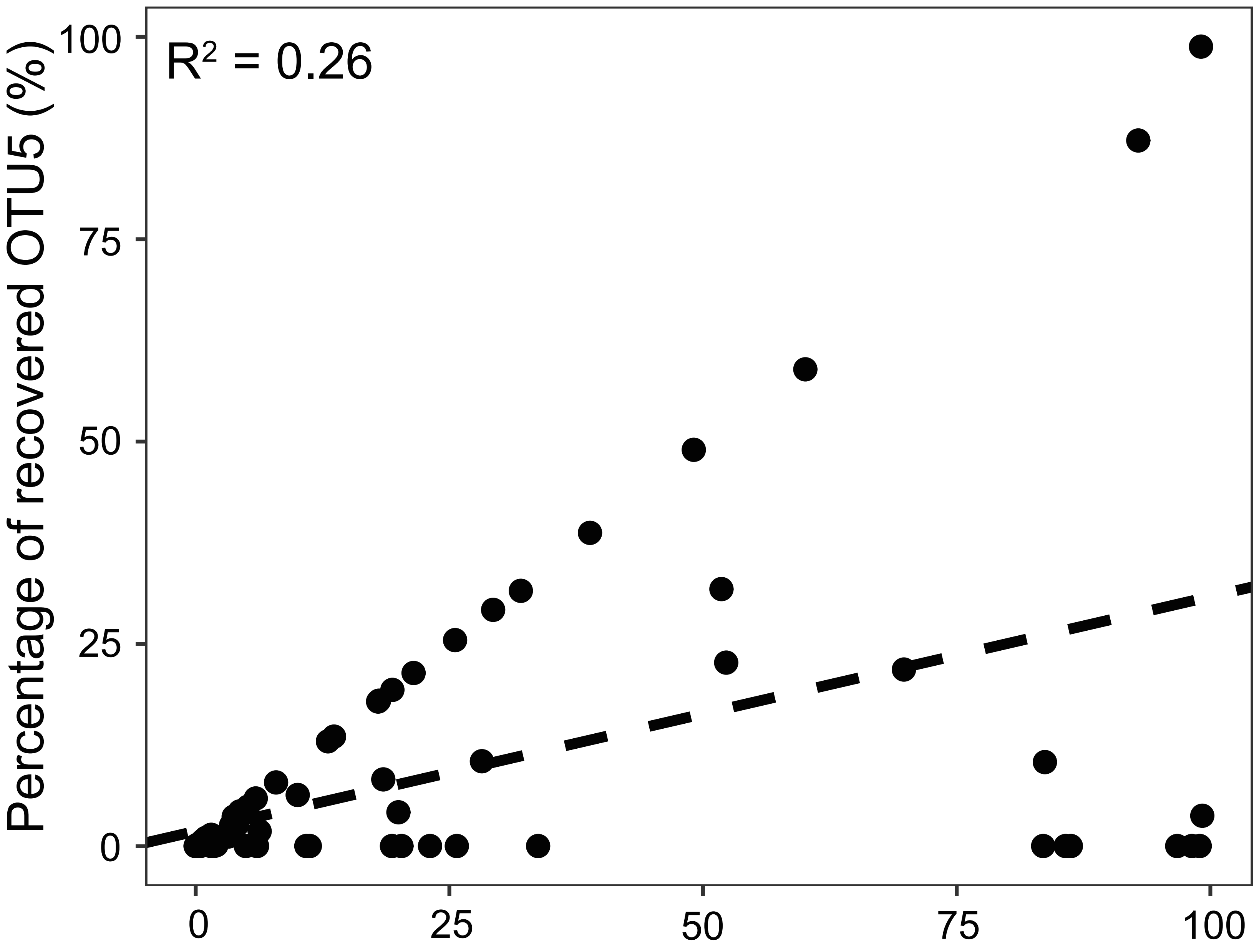
